# Meta-transcriptomic comparison of the RNA viromes of the mosquito vectors *Culex pipiens* and *Culex torrentium* in northern Europe

**DOI:** 10.1101/725788

**Authors:** John H.-O. Pettersson, Mang Shi, John-Sebastian Eden, Edward C. Holmes, Jenny C. Hesson

## Abstract

There is mounting evidence that mosquitoes harbour an extensive diversity of ‘insect-specific’ RNA viruses in addition to those important to human and animal health. However, because most studies of the mosquito virome have been conducted at lower latitudes there is a major knowledge gap on the genetic diversity, evolutionary history, and spread of RNA viruses sampled from mosquitoes in northern latitudes. Here, we determined and compared the RNA virome of two common northern *Culex* mosquito species, *Cx. pipiens* and *Cx. torrentium*, known vectors of West Nile virus and Sindbis virus, respectively, collected in south-central Sweden. Following bulk RNA-sequencing (meta-transcriptomics) of 12 libraries, comprising 120 specimens of *Cx. pipiens* and 150 specimens of *Cx. torrentium*, we identified 40 viruses (representing 14 virus families) of which 28 were novel based on phylogenetic analysis of the RNA-dependent RNA polymerase (RdRp) protein. Hence, we found similar levels of virome diversity as in mosquitoes sampled from the more biodiverse lower latitudes. Four libraries, all from *Cx. torrentium*, had a significantly higher abundance of viral reads, spanning ∼7– 36% of the total amount of reads. Many of these viruses were also related to those sampled on other continents, indicative of widespread global movement and/or long host-virus co-evolution. Importantly, although the two mosquito species investigated have overlapping geographical distributions and share many viruses, approximately one quarter of the viruses were only found at a specific location, such that geography must play an important role in shaping the diversity of RNA viruses in *Culex* mosquitoes.

**Importance:** RNA viruses are found in all domains of life and all global habitats. However, the factors that determine virome composition and structure within and between organisms are largely unknown. Herein, we characterised RNA virus diversity in two common mosquito vector species, *Culex pipiens* and *Culex torrentium*, sampled from northern Europe. Our analysis revealed extensive viral diversity, including 28 novel viruses, and was comparable to the levels of diversity found in other temperate and tropical regions globally. Importantly, as well as harbouring RNA viruses that are closely related to other mosquito-derived viruses sampled in diverse global locations, we also described a number of viruses that are unique to specific sampling locations in Sweden. Hence, these data showed that geographical factors can play an important role in shaping virome structure even at local scales.

## Introduction

The mosquito (Diptera; *Culicidae*) genus *Culex* comprises more than a thousand species, with representatives found globally (1). *Culex* species are vectors of a number of important pathogens including West Nile virus (WNV) (*Flaviviridae*), Japanese encephalitis virus (JEV) (*Flaviviridae*) and Sindbis virus (SINV) (*Togaviridae*), as well as a variety of nematodes (1–3). One of the most widespread *Culex* species is the Northern House mosquito, *Cx. pipiens*, that is distributed across the northern hemisphere. In Europe and the Middle East it occurs together with *Cx. torrentium*, another *Culex* species with females and larvae that are morphologically identical to *Cx. pipiens*. These two species have overlapping distributions and share larval habitats. However, *Cx. torrentium* dominates in northern Europe while *Cx. pipiens* is more abundant in the south (4). Both species are vectors for a number of bird-associated viruses that can cause disease in Europe; for example, WNV, that may cause a febrile disease with encephalitis, and SINV that may result in long lasting arthritis (2, 5). *Cx. pipiens* is one of the most common WNV vectors in both southern Europe and North America, and *Cx. torrentium* is the main vector of SINV in northern Europe (2, 6). Infections with these pathogenic viruses occur in late summer when the viral prevalence accumulates in passerine birds, the vertebrate hosts of both of these viruses (7, 8). Despite their importance as vectors, little is known about the detailed biology of *Cx. pipiens* and *Cx. torrentium* due to the difficulties in species identification, which can only be reliably achieved through molecular means. Much of the biology of these species, such as their larval habitat and feeding preferences, is considered similar. However, one significant difference between the two species is that while *Cx. pipiens* harbours a high prevalence of the intracellular bacteria *Wolbachia pipientis*, it is seemingly absent in *Cx. torrentium* (9).

In recent years, studies utilizing RNA-sequencing (RNA-Seq, or ‘meta-transcriptomics’) have revealed an enormous RNA virus diversity in both vertebrates and invertebrates (10, 11). Mosquitoes are of particular interest as many are well-known vectors of pathogenic viruses. Importantly, recent studies have shown that these pathogenic viruses represent only a fraction of the total virome in the mosquito species investigated. Indeed, mosquitoes clearly carry a large number of newly described and divergent arthropod-specific viruses, with representatives from many genetically diverse virus families and orders, such as the *Flaviviridae, Togaviridae* and the *Bunyavirales* (12–16). However, most studies have been conducted on latitudes below 55°, such that there is a marked lack of data of the mosquito viral diversity present in northern temperate regions where the composition of mosquito species as well as environmental parameters differ significantly from lower latitudes. Indeed, for many forms of life, biodiversity increases towards the equator (17), and the species richness of mosquitoes is greater in tropical regions than temperate regions (18). A central aim of the current study was therefore to investigate whether viral diversity co-varies in the same manner. Given that *Cx. pipiens* and *Cx. torrentium* are two common *Culex* species in northern and central Europe, and known vectors of SINV and WNV, they were chosen for RNA virome investigation and comparison by RNA-Seq.

## Results

### RNA virome characterisation

We characterised the RNA viral transcriptome of two mosquito species, *Cx. pipiens* and *Cx. torrentium*, collected from central and southern Sweden (Suppl. table 1). After high-throughput sequencing, a total of 569,518,520 (range 34,150,856–62,936,342) 150bp reads were produced from 12 ribosomal RNA-depleted sequence libraries that were assembled into 153,583 (4,333–33,893) contigs. From all the contigs assembled, we identified 40 that contained RdRp sequence motifs and hence indicative of viruses, belonging to 14 different viral families/orders: *Alphaviridae, Bunyavirales, Endornaviridae, Luteoviridae, Mononegavirales, Nidovirales, Orthomyxoviridae, Partitiviridae, Picornaviridae, Reoviridae, Totiviridae, Tymoviridae* and representatives from the divergent *Virgaviridae, Negeviridae*- and Qin-viruses. For each viral family/order, between one and five virus species were identified and in total 28 novel RNA virus species were discovered here, which were named based on geographical location.

The relative number of all virus reads, as mapped to contigs with RdRp-motifs, compared to the total amount of non-ribosomal RNA reads per library varied between 0.1–36.6% (Table 1). Notably, libraries 2, 10, 11 and 12 from *Cx. torrentium* were characterised by a significantly higher number of viral reads compared to non-viral reads (Figure 1, Table 1). The individual abundance of each viral species, measured as the number of reads mapped to each RdRp contig divided by the total amount of reads in the library × 1000,000 (i.e. reads per million, RPM), varied between 1.09–10,006.67 RPM for *Cx. pipiens* and 1.08–303,145.83 RPM for *Cx. torrentium*. In comparison, the abundance of host reads, as measured by the presence of the host mitochondrial protein COX1, was more stable and varied only between 4.22–66.99 RPM across all libraries (Table 2).

**Table 1.** Overview of viral RdRp-motif library content compared to the total number of non-ribosomal RNA reads per library.

| Library | *Cx torrentium* |  |  |  |  |  | *Cx pipiens* |  |  |  |  |  |
| --- | --- | --- | --- | --- | --- | --- | --- | --- | --- | --- | --- | --- |
|  | L1 | L2 | L9 | L10 | L11 | L12 | L3 | L4 | L5 | L6 | L7 | L8 |
| **Total virus reads** | 311,893 | 22,961,076 | 258,016 | 9,604,141 | 4,654,540 | 7,452,859 | 45,019 | 279,568 | 565,968 | 57,867 | 227,988 | 186,792 |
| **Host COX1 reads** | 586 | 265 | 322 | 2427 | 2254 | 3117 | 126 | 2529 | 317 | 2850 | 860 | 417 |
| **Total reads** | 34,150,856 | 62,820,620 | 43,914,132 | 52,916,282 | 62,936,342 | 59,016,596 | 39,231,440 | 41,210,662 | 41,328,330 | 40,762,624 | 44,703,752 | 46,526,884 |
| **Virus %** | 0.9133 | 36.5502 | 0.5875 | 18.1497 | 7.3956 | 12.6284 | 0.1148 | 0.6784 | 1.3694 | 0.1420 | 0.5100 | 0.4015 |
| **Host COX1 %** | 0.0017 | 0.0004 | 0.0007 | 0.0046 | 0.0036 | 0.0053 | 0.0003 | 0.0061 | 0.0008 | 0.0070 | ..0019 | ..0009 |
| **Other %** | 99.0850 | 63.4494 | 99.4117 | 81.8457 | 92.6008 | 87.3663 | 99.8849 | 99.3155 | 98.6298 | 99.8510 | 99.4881 | 99.5976 |

**Table 2.** Individual abundance, measured as reads per million, of each virus, as well as *Wolbachia* bacteria, in comparison with the abundance of host COX1 gene.

|  |  |  |  | Cx torrentium |  |  |  |  |  | Cx pipiens |  |  |  |  |  |
| --- | --- | --- | --- | --- | --- | --- | --- | --- | --- | --- | --- | --- | --- | --- | --- |
|  |  |  | Location | SINV+ |  | SINV unscreened |  |  |  | SINV+ | SINV unscreened |  |  |  |  |
| Virus | Threshold |  | # mosq | Dalälven | Dalälven | Dalälven | Dalälven | Dalälven | Dalälven | Dalälven | Dalälven | Dalälven | Kristianstad | Kristianstad | Dalälven |
|  | * | ** |  | 10 | 10 | 50 | 50 | 15 | 15 | 10 | 15 | 15 | 15 | 15 | 50 |
|  |  |  | Lenght | L1 | L2 | L9 | L10 | L11 | L12 | L3 | L4 | L5 | L6 | L7 | L8 |
| Sindbis virus | 0,672 | 1 | 11,688 | 671,784 | 417,443 | 0,387 | 0,151 | 0,540 | 0,474 | 2,523 | 0,995 | 0,339 | 2,012 | 0,380 | 0,107 |
| Nam Dinh virus | 303,146 | 1 | 20,240 | 73,673 | 303145,830 | 85,644 | 120821,017 | 28690,339 | 95,244 | 137,084 | 83,376 | 10006,671 | 116,626 | 87,174 | 106,154 |
| Biggie virus | 35,064 | 1 | 9207 | 16,720 | 15525,826 | 15,439 | 34543,772 | 23,405 | 35063,781 | 46,391 | 18,709 | 20,978 | 18,301 | 21,363 | 28,930 |
| Negev virus | 9,715 | 1 | 9493 | 1,171 | 9715,234 | 1,070 | 2,192 | 1,748 | 1,915 | 2,651 | 1,432 | 1,670 | 2,723 | 1,946 | 582,029 |
| Rinkaby virus | 0,706 | 1 | 14,498 | 0,000 | 0,159 | 0,068 | 0,151 | 0,127 | 0,034 | 0,280 | 0,000 | 0,387 | 0,196 | 706,361 | 0,172 |
| Kerstinbo virus | 1,825 | 1 | 11,280 | 0,381 | 0,207 | 0,250 | 10,734 | 117,913 | 0,136 | 156,456 | 1825,401 | 0,266 | 0,098 | 146,475 | 20,633 |
| Forneby virus | 0,138 | 1 | 8255 | 0,000 | 0,064 | 0,000 | 3,288 | 0,000 | 0,000 | 0,051 | 138,241 | 0,000 | 0,049 | 0,000 | 6,362 |
| Osterfarnebo virus | 0,087 | 1 | 5357 | 0,000 | 0,000 | 0,046 | 1,077 | 0,000 | 0,000 | 0,000 | 87,016 | 0,000 | 0,000 | 0,000 | 4,900 |
| Hallsjon virus | 0,014 | 1 | 2847 | 0,000 | 0,000 | 0,000 | 0,491 | 13,712 | 0,000 | 0,000 | 0,340 | 0,048 | 0,442 | 0,000 | 0,000 |
| Tarnsjo virus (variant 1) | 0,063 | 1 | 7890 | 49,047 | 0,000 | 11,955 | 27,629 | 63,064 | 2,864 | 0,051 | 0,000 | 0,024 | 14,253 | 0,045 | 0,086 |
| Tarnsjo virus (variant 2) | 0,076 | 1 | 7838 | 7,701 | 0,000 | 2,095 | 3,553 | 10,471 | 0,491 | 0,000 | 0,000 | 0,048 | 76,075 | 0,000 | 0,021 |
| Culex associated luteo like virus | 1,101 | 1 | 2790 | 0,322 | 0,127 | 0,182 | 0,113 | 0,127 | 0,322 | 0,510 | 0,170 | 0,387 | 0,368 | 1101,496 | 109,421 |
| Berrek virus | 0,014 | 1 | 2807 | 0,000 | 0,000 | 0,046 | 0,000 | 0,000 | 0,034 | 0,000 | 0,000 | 0,000 | 0,000 | 0,000 | 13,906 |
| Fagle virus | 0,002 | 1 | 1453 | 0,000 | 1,544 | 0,000 | 0,000 | 0,000 | 0,000 | 0,000 | 0,000 | 0,000 | 0,000 | 0,000 | 0,000 |
| Marma virus | 2,602 | 1 | 3151 | 0,703 | 0,493 | 0,934 | 1,077 | 0,604 | 0,729 | 1,963 | 0,849 | 0,919 | 1,251 | 1737,371 | 2601,593 |
| Merida virus | 12,423 | 1 | 11,785 | 2327,789 | 2408,604 | 2230,876 | 7176,373 | 7,420 | 12423,082 | 16,084 | 9,803 | 2037,222 | 6,526 | 6,219 | 301,911 |
| Culex mononega like virus 2 | 1,216 | 1 | 13,316 | 467,748 | 157,082 | 285,831 | 1216,374 | 553,242 | 311,692 | 1,096 | 467,088 | 87,857 | 13,296 | 0,761 | 23,513 |
| Gysinge virus | 0,260 | 1 | 9532 | 0,088 | 0,032 | 0,091 | 0,170 | 7,245 | 0,102 | 115,749 | 259,544 | 0,073 | 0,147 | 68,093 | 0,086 |
| Culex mosquito virus 4 | 0,453 | 1 | 11,954 | 0,059 | 452,511 | 0,068 | 1,928 | 0,079 | 1,881 | 0,102 | 0,000 | 0,097 | 0,270 | 0,045 | 0,129 |
| Culex mononega like virus 1 | 0,083 | 1 | 6604 | 0,000 | 0,000 | 0,046 | 0,038 | 0,000 | 0,000 | 0,000 | 0,000 | 0,000 | 0,000 | 0,000 | 83,113 |
| Valmbacken virus | 52,265 | 1 | 4315 | 19,092 | 32360,489 | 139,363 | 1158,301 | 29,172 | 52264,756 | 66,401 | 24,678 | 21,462 | 19,331 | 20,983 | 30,993 |
| Jotan virus | 20,309 | 1 | 9112 | 16,808 | 18,433 | 15,712 | 7758,198 | 41738,015 | 20308,796 | 50,878 | 91,457 | 912,788 | 18,718 | 18,119 | 22,224 |
| Ista virus | 5,605 | 1 | 9551 | 5605,247 | 576,435 | 2529,573 | 7361,250 | 2402,094 | 3661,343 | 8,131 | 8,129 | 4,065 | 3,950 | 4,094 | 116,771 |
| Wuhan Mosquito Virus 4 | 1,900 | 1 | 2445 | 1,318 | 229,558 | 520,561 | 642,373 | 332,654 | 1679,358 | 677,569 | 1900,406 | 1,742 | 325,740 | 930,772 | 70,003 |
| Wuhan Mosquito Virus 6 | **0,636** | **1** | 2440 | 0,117 | 0,127 | 0,774 | 0,189 | 0,127 | 0,102 | 0,102 | 0,097 | 636,319 | 510,664 | 2,349 | 1,891 |
| Vivastbo virus | **1,338** | **1** | 2157 | 0,000 | 0,096 | 0,091 | 34,791 | 3,972 | 0,169 | 7,341 | 1338,440 | 0,266 | 0,393 | 119,475 | 26,974 |
| Sonnbo virus | **0,088** | **1** | 1737 | 0,000 | 0,032 | 0,000 | 0,038 | 0,064 | 0,034 | 0,000 | 87,720 | 0,048 | 0,000 | 0,000 | 15,561 |
| Rasbo virus | **0,017** | **1** | 5974 | 0,000 | 0,000 | 4,964 | 7,257 | 4,798 | 17,368 | 0,051 | 0,049 | 0,000 | 0,000 | 0,000 | 0,043 |
| Kristianstad virus | **0,022** | **1** | 5406 | 0,000 | 0,000 | 0,000 | 0,038 | 0,000 | 0,000 | 0,000 | 0,000 | 0,000 | 22,104 | 0,000 | 0,000 |
| Asum virus | **0,362** | **1** | 7184 | 0,000 | 0,032 | 0,410 | 0,397 | 0,016 | 0,051 | 0,051 | 0,218 | 0,097 | 361,949 | 0,045 | 0,043 |
| Salari virus | **0,014** | **1** | 6630 | 0,000 | 0,000 | 0,000 | 0,000 | 0,000 | 0,000 | 0,051 | 0,000 | 0,000 | 0,000 | 0,045 | 13,906 |
| Anjon virus | **0,407** | **1** | 6495 | 3,485 | 406,952 | 140,775 | 725,051 | 6,864 | 537,290 | 0,918 | 0,582 | 1,089 | 0,589 | 0,224 | 0,645 |
| Gran virus | **0,012** | **1** | 5622 | 0,000 | 0,032 | 9,473 | 5,858 | 3,559 | 11,912 | 0,000 | 0,000 | 0,048 | 0,000 | 0,000 | 0,021 |
| Nackenback virus | **0,559** | **1** | 6128 | 0,059 | 0,064 | 0,091 | 0,113 | 8,374 | 0,102 | 0,102 | 559,443 | 0,048 | 0,123 | 65,498 | 0,107 |
| Vinslov virus | **0,047** | **1** | 5590 | 0,000 | 0,000 | 0,046 | 0,000 | 0,000 | 0,000 | 0,000 | 0,340 | 8,082 | 47,445 | 0,000 | 0,000 |
| Vittskovle virus | **0,019** | **1** | 5671 | 0,000 | 0,684 | 0,046 | 0,000 | 0,000 | 0,000 | 0,000 | 0,000 | 0,000 | 19,380 | 0,000 | 0,000 |
| Ahus virus | **0,027** | **1** | 7732 | 0,000 | 0,032 | 0,000 | 0,038 | 0,000 | 0,000 | 0,000 | 0,243 | 4,404 | 26,691 | 0,000 | 0,000 |
| Osta virus | **0,019** | **1** | 5398 | 0,000 | 0,032 | 0,000 | 0,000 | 0,000 | 0,000 | 0,102 | 19,364 | 0,000 | 0,049 | 10,111 | 0,043 |
| Lindangsbacken virus | **0,105** | **1** | 6171 | 0,000 | 104,711 | 0,046 | 0,000 | 0,000 | 0,000 | 0,000 | 0,000 | 0,000 | 0,000 | 0,045 | 0,000 |
| Salja virus | **0,002** | **1** | 1286 | 0,000 | 0,000 | 0,000 | 0,000 | 0,000 | 0,000 | 0,000 | 1,626 | 0,000 | 0,000 | 1,611 | 0,000 |
| Eskilstorp virus | **0,210** | **1** | 2933 | 0,000 | 0,000 | 0,000 | 0,076 | 0,032 | 0,068 | 0,076 | 0,000 | 0,000 | 0,049 | 210,363 | 0,129 |
| Wolbachia COX1 | **0,001** | **1** | 1573 | 0,00 | 0,99 | 0,00 | 0,00 | 0,00 | 0,00 | 0,00 | 0,05 | 0,05 | 0,00 | 0,00 | 0,00 |
| Wolbachia WSP | **0,000** | **1** | 614 | 0,00 | 0,00 | 0,00 | 0,00 | 0,03 | 0,20 | 0,10 | 0,34 | 0,22 | 0,00 | 0,00 | 0,09 |
| Host COX1 | **0,067** | **1** | 1506 | 17,16 | 4,22 | 7,33 | 45,86 | 35,81 | 52,82 | 14,94 | 6,43 | 7,79 | 59,54 | 50,42 | 66,99 |
| Number of virus species | — | — | — | 6 | 13 | 9 | 17 | 14 | 12 | 10 | 13 | 8 | 10 | 12 | 17 |

\* = 0,1% of the most abundant library
\*\* = Less than 1 per million readds
Grey = not present
Green = present, but not abundant
Orange = present and >0.1% of total reads
Red = present and more abundant than host

**Figure 1.**
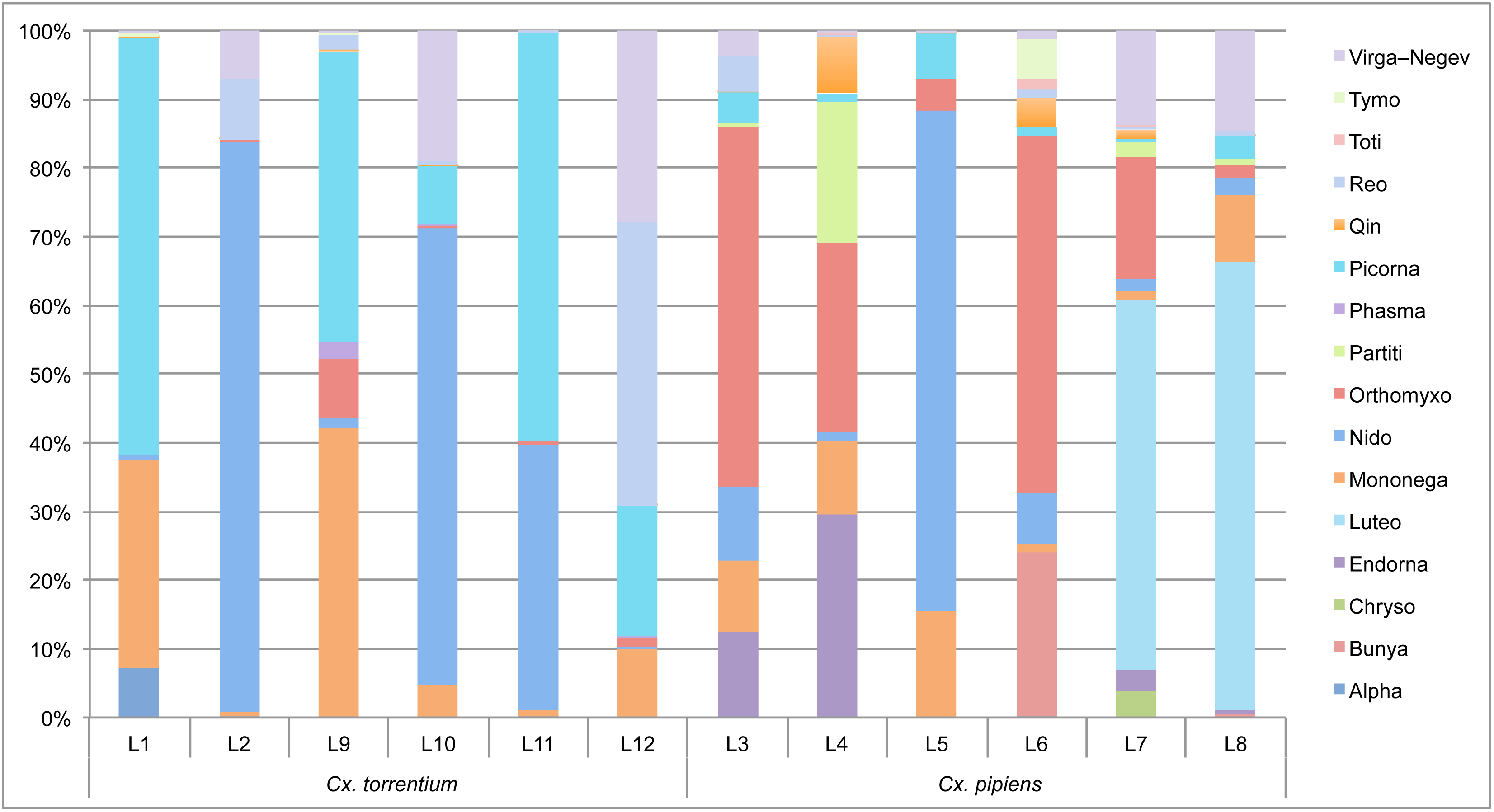
Estimation of virome composition compared to host (non-viral) content in each library.

### Virome comparison between mosquito species and geographical regions

Both the composition and abundance of the virus species and families observed differs between the two mosquito species (Figure 2, Table 2). Of the 40 virus species discovered, most were found in *Cx. pipiens* which harboured 34 species: 23 of these are newly described in *Cx. pipiens* and 11 have been described previously. Sixteen of these 34 virus species were unique to *Cx. pipiens* and hence not present in *Cx. torrentium*. Similarly, 24 of the 40 virus species were discovered in *Cx. torrentium*: 18 of these are newly described in *Cx. torrentium* and 6 have been described previously. Six viruses found *Cx. torrentium* were not present in *Cx. pipiens* (Figure 3, Table 2).

**Figure 2.**
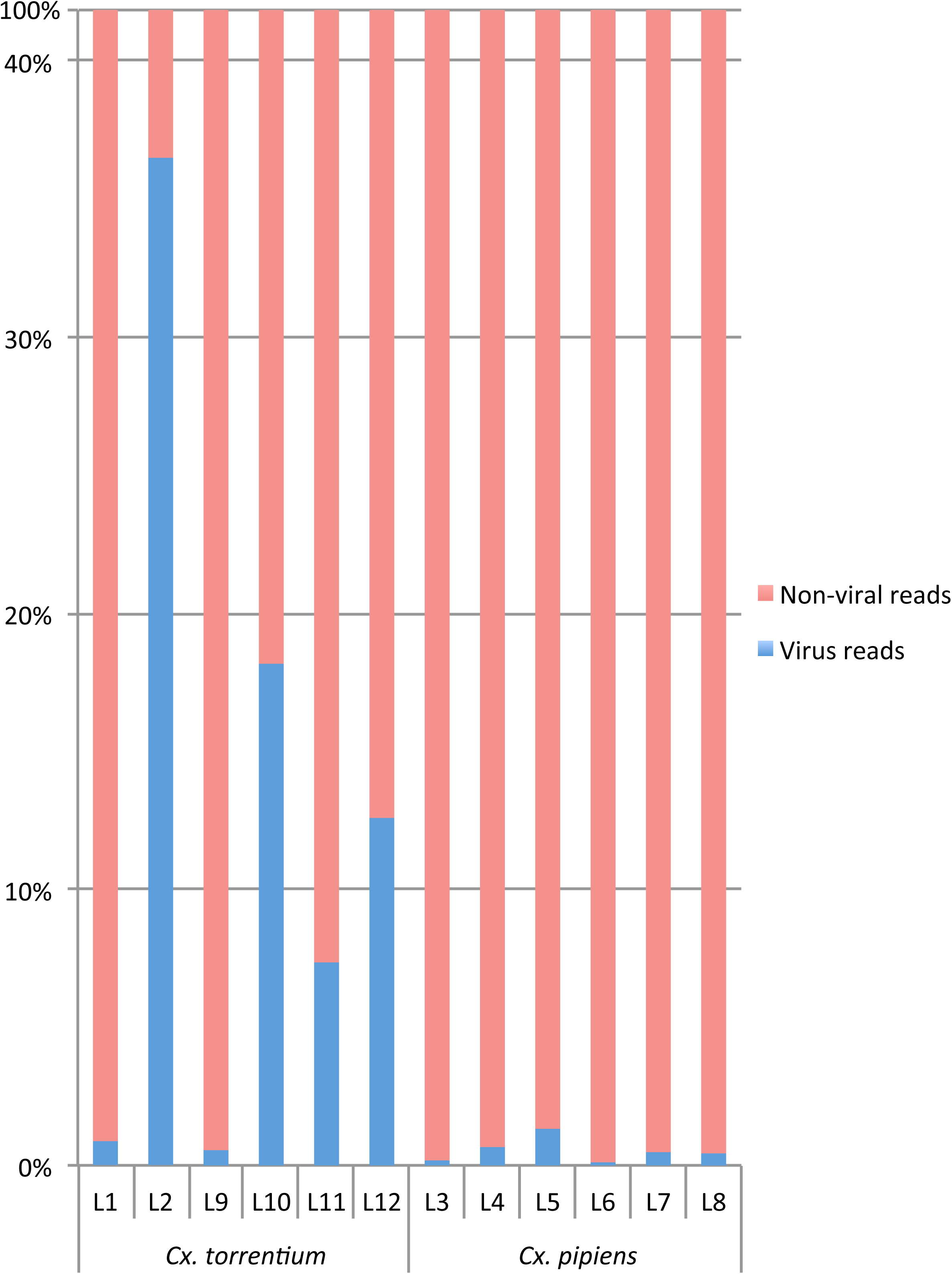
Comparison of the virome family composition and abundance between the *Cx. pipiens* and *Cx. torrentium.* For ease of presentation, abbreviations are used to indicate virus taxonomy in each case.

**Figure 3.**
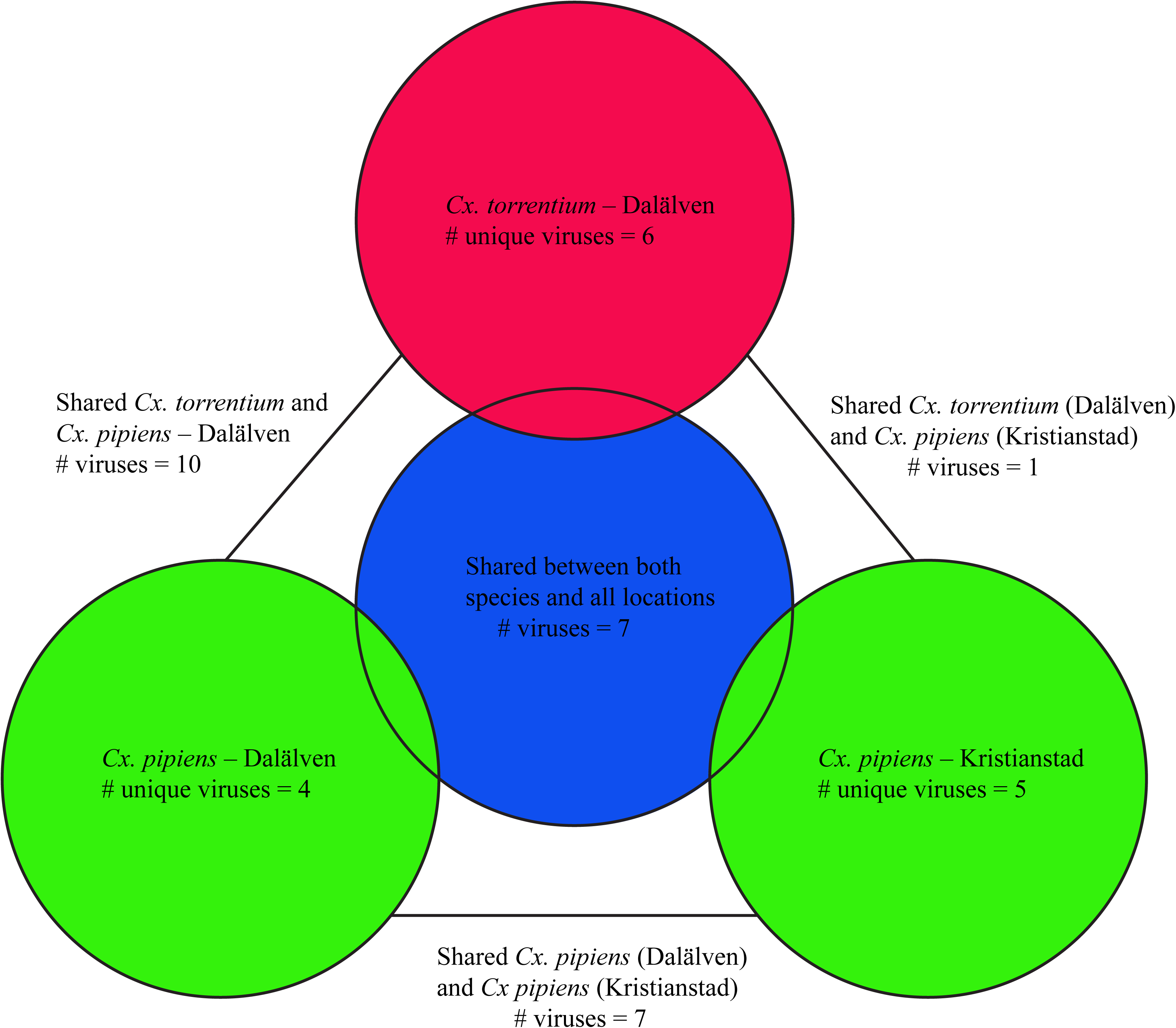
Venn diagram showing the number of unique and shared viruses per location per mosquito species.

We next analysed potential host-relationships by comparing abundance, presence across libraries and phylogenetic relationship with other viruses (Table 3). These data suggest that 16 of the 40 viruses were likely mosquito associated, of which one and two were unique to *Cx. pipiens* and *Cx. torrentium*, respectively (Figure 4–6). The host association was unclear in the remaining viruses (for example, they could be associated with micro-organisms co-infecting the mosquitoes) and could not be safely assumed to infect mosquitoes. For example, Ahus virus (*Totiviridae*) was at low abundance, was not present in several libraries, and clustered with viruses derived from various environmental samples, suggesting that it is less likely to be mosquito-associated. Similarly, although Gysinge virus (*Mononegavirales*) was abundant and present in several libraries (Table 3), that its closest relative (Figure 5) was a soy bean leaf associated virus (19) means that it cannot be safely assigned to mosquitoes. Conversely, Culex mononega like virus 2 (Figure 5) was found to be abundant, present in several libraries and clustered with other mosquito viruses, suggesting that it is likely to be mosquito associated. All potential mosquito host-association data is summarised in Table 3.

**Table 3.** Indication of host associations for the discovered viruses. Host-association was assessed using (i) the abundance of viral contig per total amount of reads in a library, (ii) virus abundance in relation to the COX1 host gene, (iii) presence amongst the libraries, and (iv) phylogenetic clustering with other mosquito derived viruses.

| Virus | Virus family | *Cx torrentium* | *Cx pipiens* | Abundant? | More abundant than host RNA? | Present in >2 libraries? | Clusters with mosquito viruses? | Mosquito associated? |
| --- | --- | --- | --- | --- | --- | --- | --- | --- |
| Sindbis virus | Alpha | P | P | Yes | Yes | No | Yes | Yes |
| Nam Dinh virus | Nido | P | P | Yes | Yes | Yes | Yes | Yes |
| Biggie virus | Virga-Negev | P | P | Yes | Yes | Yes | Yes | Yes |
| Negev virus | Virga-Negev | P | P | Yes | Yes | No | Yes | Yes |
| Rinkaby virus | Virga-Negev | NP | P | Yes | Yes | No | No | Yes |
| Kerstinbo virus | Endorna | P | P | Yes | Yes | No | No | Yes |
| Forneby virus | Endorna | P | P | No | Yes | No | No | No |
| Osterfarnebo virus | Endorna | P | P | No | Yes | No | No | No |
| Hallsjon virus | Endorna | P | NP | No | No | No | No | No |
| Tarnsjo virus (variant 1) | Tymo | P | P | No | Yes | No | Yes | Yes |
| Tarnsjo virus (variant 2) | Tymo | P | P | No | Yes | No | Yes | Yes |
| Culex associated luteo like virus | Luteo | NP | P | Yes | Yes | No | Yes | Yes |
| Berrek virus | Luteo | NP | P | No | No | No | Yes | No |
| Fagle virus | Luteo | P | NP | No | No | No | No | No |
| Marma virus | Luteo | NP | P | Yes | Yes | No | Yes | Yes |
| Merida virus | Mononega | P | P | Yes | Yes | Yes | Yes | Yes |
| Culex mononega like virus 2 | Mononega | P | P | Yes | Yes | Yes | Yes | Yes |
| Gysinge virus | Mononega | P | P | Yes | Yes | Yes | No | Yes |
| Culex mosquito virus 4 | Mononega | P | NP | Yes | Yes | No | Yes | Yes |
| Culex mononega like virus 1 | Mononega | NP | P | No | Yes | No | Yes | Yes |
| Valmbacken virus | Reo | P | P | Yes | Yes | Yes | Yes | Yes |
| Jotan virus | Picornia | P | P | Yes | Yes | Yes | Yes | Yes |
| Ista virus | Picornia | P | P | Yes | Yes | Yes | No | Yes |
| Wuhan Mosquito Virus 6 | Orthomyxo | P | P | Yes | Yes | Yes | Yes | Yes |
| Wuhan Mosquito Virus 4 | Orthomyxo | NP | P | Yes | No | No | Yes | Yes |
| Vivastbo virus | Partiti | P | P | Yes | Yes | No | No | Yes |
| Sonnbo virus | Partiti | NP | P | No | Yes | No | No | No |
| Rasbo virus | Bunya | P | NP | No | No | No | No | No |
| Kristianstad virus | Bunya | NP | P | No | No | No | Yes | No |
| Asum virus | Bunya | NP | P | Yes | No | No | No | No |
| Salari virus | Bunya | NP | P | No | No | No | Yes | No |
| Anjon virus | Phasma | P | P | Yes | Yes | Yes | Yes | Yes |
| Gran virus | Qin | P | NP | No | No | No | No | No |
| Nackenback virus | Qin | P | P | Yes | Yes | Yes | No | Yes |
| Vinslov virus | Qin | NP | P | No | No | No | No | No |
| Vittskovle virus | Qin | NP | P | No | No | No | No | No |
| Ahus virus | Toti | NP | P | No | No | No | No | No |
| Osta virus | Toti | NP | P | No | No | No | Yes | No |
| Lindangsbacken virus | Toti | P | NP | No | Yes | No | Yes | Yes |
| Salja virus | Toti | NP | P | No | No | No | Yes | No |
| Eskilstorp virus | Chryso | NP | P | No | Yes | No | Yes | **Yes** |

P = Present NP = Not present
>0.1% = abundant

**Figure 4.**
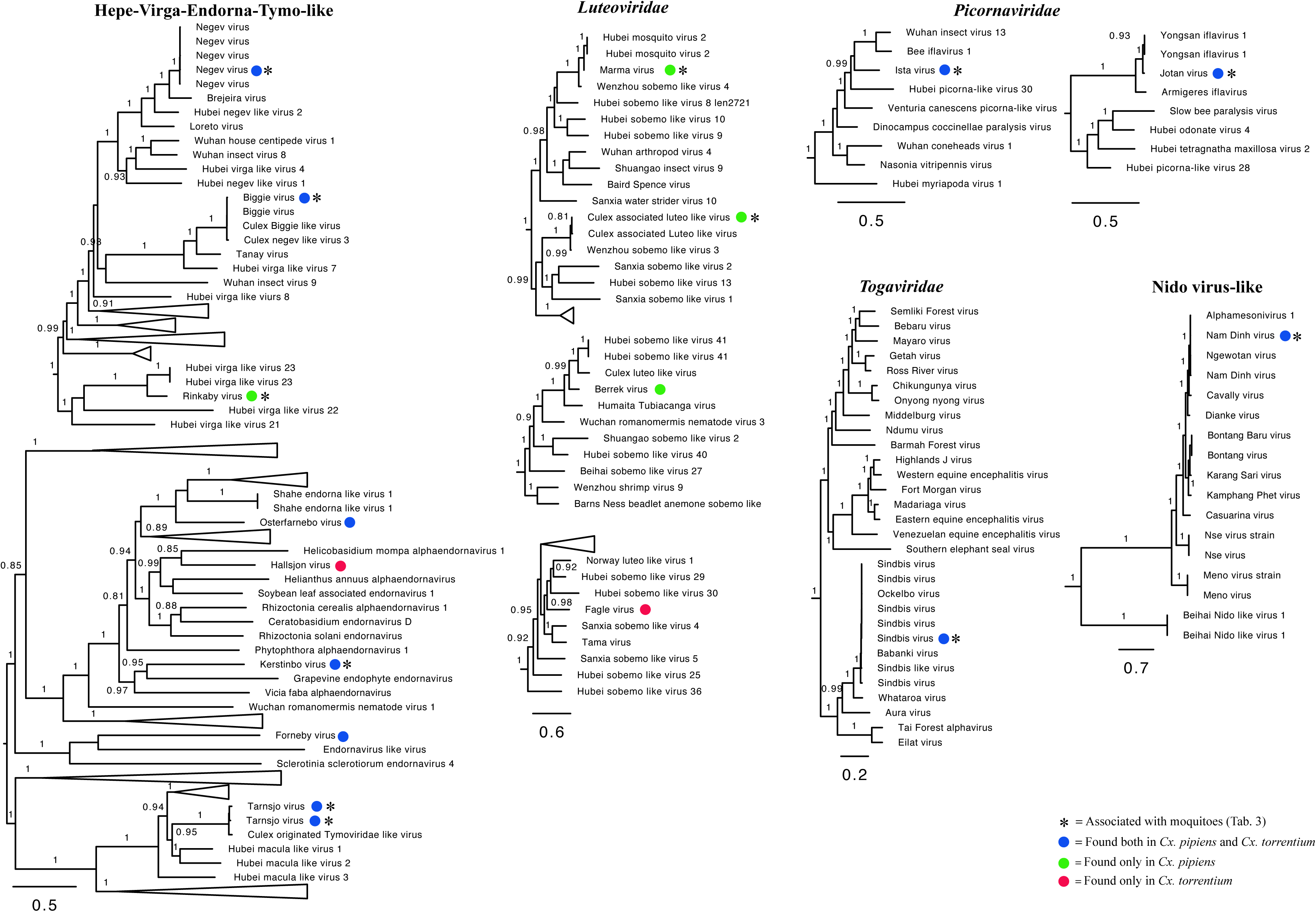
Phylogenetic analysis of all the positive-sense RNA viruses identified here (marked by coloured circles) along with representative publicly available viruses. Those viruses most likely associated with mosquitoes are marked by an *. Numbers on branches indicate SH support, and only branches with SH support ≥80% are indicated. Branch lengths are scaled according to the number of amino acid substitutions per site. All phylogenetic trees were midpoint-rooted for clarity only.

**Figure 5.**
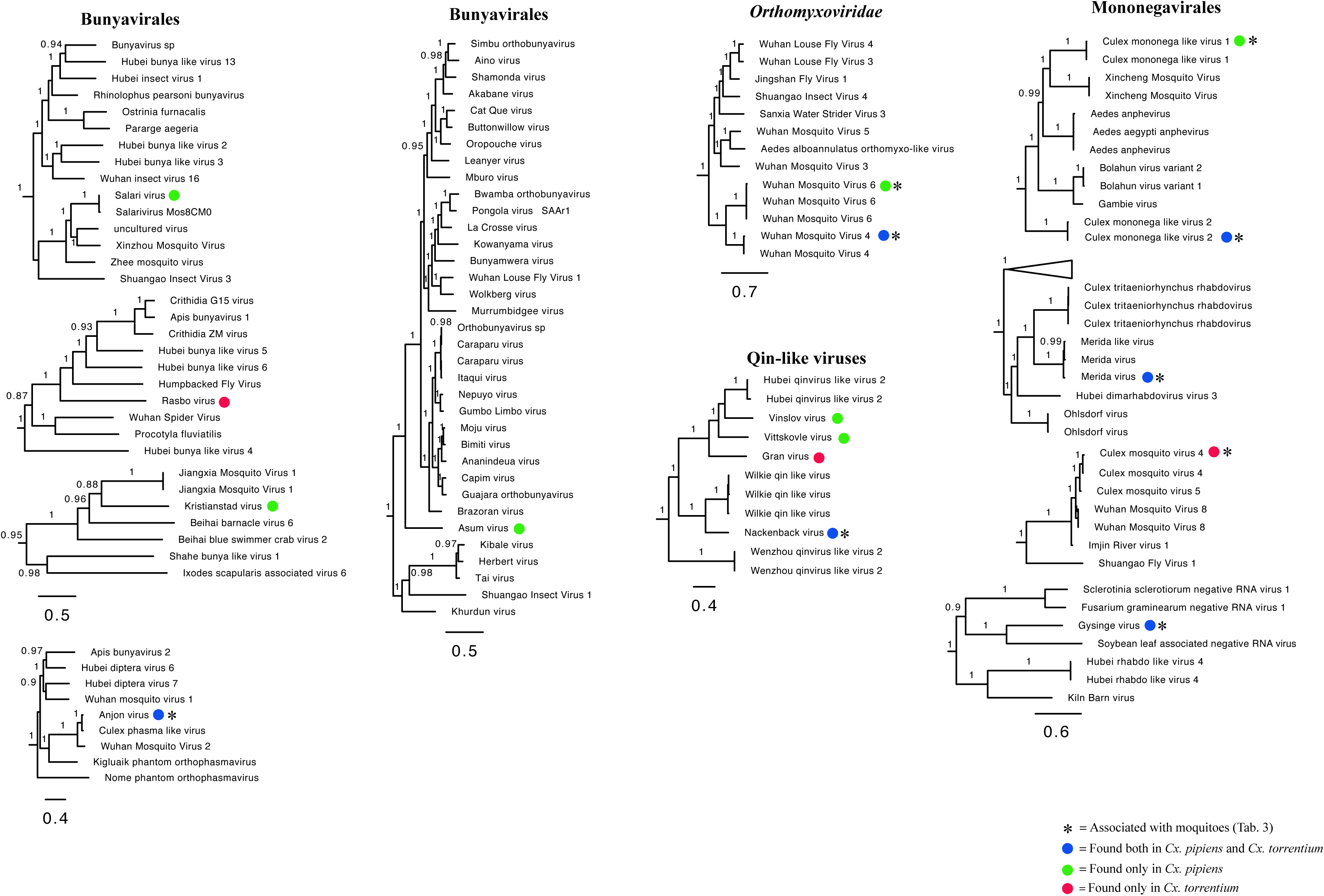
Phylogenetic analysis of all the negative-sense RNA viruses identified here (marked by coloured circles) along with representative publicly available viruses. Those viruses most likely associated with mosquitoes are marked by an *. Numbers on branches indicate SH support, and only branches with SH support ≥80% are indicated. Branch lengths are scaled according to the number of amino acid substitutions per site. All phylogenetic trees were midpoint-rooted for clarity only.

**Figure 6.**
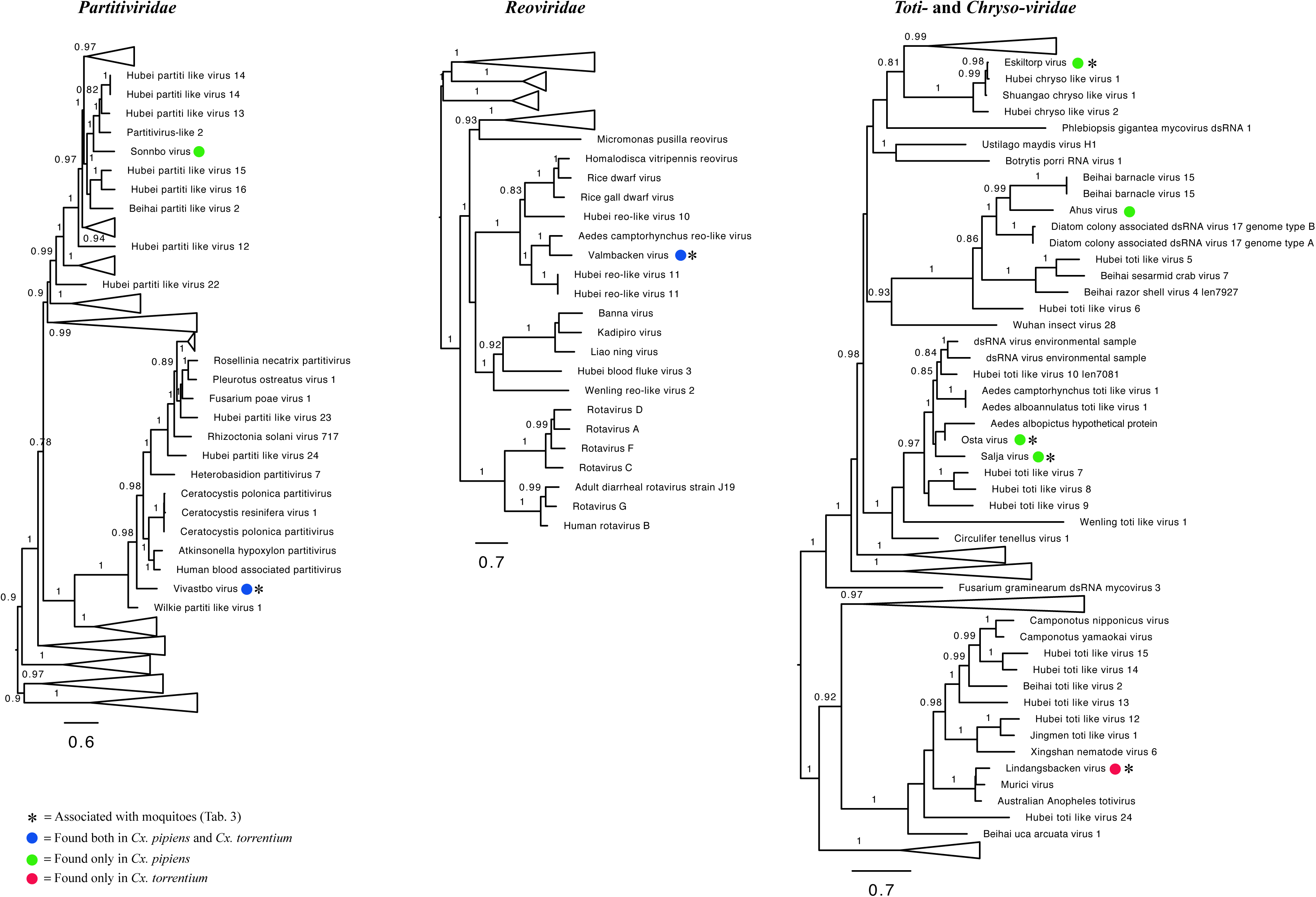
Phylogenetic analysis of all the double-stranded RNA viruses identified here (marked by coloured circles) along with representative publicly available viruses. Those viruses most likely associated with mosquitoes are marked by an *. Numbers on branches indicate SH support, and only branches with SH support ≥80% are indicated. Branch lengths are scaled according to the number of amino acid substitutions per site. All phylogenetic trees were midpoint-rooted for clarity only.

Notably, *Cx. torrentium* harboured four viruses of significantly higher abundance compared to *Cx. pipiens*: (i) Nam Dinh virus (303,145 RPM, or 42% of all viral reads and more than 30% of all [non rRNA] reads, respectively, in library L2), (ii) Biggie virus (35,063 RPM, or 4.6% of all viral reads and 3.5% of all reads, respectively, in library L12), as well as two newly identified viruses, (iii) Valmbacken virus (52,264 RPM, or 27% of all viral reads and 3.5% of all reads, respectively, in library L12), and (iv) Jotan virus (41,738 RPM, or 56% of all viral reads and 4.2% of all reads, respectively, in library L11) (Table 2, Suppl. table 2). *Cx. pipiens* had a more even composition of viral families across libraries (Figure 2), and the most abundant virus, Nam Dinh virus, reached 10,006 RPM (or 73% of all viral reads and 1% of all reads, respectively) in library L5.

We next compared the virome composition between Kristianstad in the south and the floodplains of the Dalälven river situated roughly 600 km further north. In the case of *Cx. pipiens* this analysis revealed a total of 20 virus species from Kristianstad, 12 of which were unique to *Cx. pipiens* and five detected in Kristianstad only, all of which were unique to *Cx. pipiens*: Asum virus (*Bunyaviridae*), Eskilstorp virus (Chrysoviridae), Kristianstad virus (*Bunyaviridae*), Rinkaby virus (Virga–Negev virus), and Vittskovle virus (*Qinvirus*). A total of 28 viruses were found in *Cx. pipiens* from Dalälven. Eleven of these were unique to *Cx. pipiens* and four were unique to *Cx. pipiens* from Dalälven: Salari virus (*Bunyavirales*), Sonnbo virus (*Partitiviridae*), Culex mononega-like virus 1 (*Mononegavirales*), and Berrek virus (*Luteoviridae*) (Table 2, Figure 2, Figure 4–6). A similar relationship was found for *Cx. torrentium*. In the case of Dalälven, 24 viruses were found in *Cx. torrentium*, of which 18 were shared with *Cx. pipiens* and six of which were unique to *Cx. torrentium* (Table 2, Figure 2, Figure 4–6). Hence, the majority of the mosquito viruses identified here were shared both between species and geographical regions, even though only 30 specimens of *Cx. pipiens* were available from Kristianstad. In contrast, approximately one quarter of the total number of virus species found at each location were unique to that location, indicative of some virome differentiation at a local geographic scale.

### Evolutionary history and host-associations of the discovered RNA viruses

Our phylogenetic analyses of the viruses discovered showed that several are closely related to previously identified viruses, and that many form clusters with mosquito-associated and/or *Culex* associated viruses within particular viral families, such as Merida virus and Gysinge virus (*Mononegavirales*), and Tarnsjo virus (*Tymovirales*) (Figure 4–6). In contrast, other novel viruses clustered with viruses neither associated with mosquitoes nor other arthropods: that they are distinguished by long branches suggests that they might infect diverse host taxa.

### Positive-sense RNA viruses

We identified 16 positive-sense RNA viruses, of which 12 were likely novel. The majority of the positive-sense RNA viruses fell within the Hepe-Virga-Endorna-Tymo-like virus complex (N = 8), whereas the others fell within *Nidovirales* (N = 1), *Luteoviridae* (N = 4), *Picornavirales* (N = 2) and *Togaviridae* (N = 1), respectively (Figure 4). The viruses discovered contain both those that are closely related to other mosquito-associated viruses, such as the highly abundant Nam Dinh virus (*Nidovirales*), as well as those without clear host associations. For example, Biggie virus clusters in a distinct group of Biggie viruses (*Virga/Endorna-viridae*) sampled from other *Culex* mosquitoes (15). We also identified several novel and divergent viruses in the family *Endornaviridae* – specifically Kerstinbo virus and Hallsjon virus – that do not cluster with other arthropod-associated viruses (Figure 4). Similarly, within the *Tymoviridae* we detected a two variants of a *Culex*-associated virus, Tarnsjo virus, that are closely related to a *Culex* associated Tymoviridae-like virus (15).

We identified four viruses within the *Luteoviridae*: *Culex* associated luteo-like virus, as well as the novel Berrek, Fagle and Marma viruses. *Culex* associated luteo like-virus has previously been found in a pool of *Culex* sp. mosquitoes from North America (15). Both of the newly discovered Berrek virus and Marma virus grouped with other luteoviruses found in mosquitoes (Figure 4), but only Marma virus was abundant, suggesting that it is *Culex* associated (Table 2, Table 3).

Two novel picornaviruses were also identified. The abundant Rinkaby virus clusters with Yongsan iflavirus 1 virus, sampled from *Cx. pipiens* mosquitoes from South Korea, and was therefore considered a *bona fide Culex* associated picornavirus. Although Ista virus did not cluster with viruses derived from mosquitoes, its high abundance and the fact that it was present in all libraries (Figure 4, Table 3) suggest that it is also *Culex* associated.

Finally, four of our libraries – L1 and L2 for *Cx. torrentium* and L3 and L6 for *Cx. pipiens* – contained reads for SINV. Importantly, whereas as the presence of SINV could be confirmed with PCR in library L1, L2 and L3, it was not PCR confirmed in L6 such that contamination cannot be excluded in this case.

### Negative-sense RNA viruses

In total, we identified 16 negative-sense RNA viruses, including 9 novel viruses. These were distributed as follows: *Bunyavirales* (N = 5), *Mononegavirales* (N = 5), Qin-like viruses (N = 4) and *Orthomyxoviridae* (N = 2) (Figure 5). As was the case for the positive-sense RNA viruses, some of these viruses that have been identified previously and cluster with viruses found in mosquitoes of the same genera, including Salari virus (*Bunyavirales*) and a number of novel viruses such as Anjon virus (*Bunyavirales*).

Within the order *Mononegavirales*, Merida virus, *Culex* mononega-like virus 2, *Culex* mosquito virus 4, and *Culex* mononega-like virus 1 have previously described been in mosquitoes (12, 15, 20). Although abundant, the novel Gysinge virus was not found to cluster with any mosquito sequences (Figure 5), so that its true host is uncertain.

The Qinviruses are a newly described and highly divergent group of RNA viruses (10). We identified four novel Qin-like viruses: Nackenback virus, Gran virus, Vinslov virus and Vittskovle virus. The latter three are more closely related to the Hubei qinvirus like virus 2 previously found in a pool of different arthropods (10). Nackenback virus was found to share a more recent common ancestor with Wilkie Qin-like virus previously found in *Aedes* and *Culex* mosquitoes in Australia (12). Although Qin-like was most closely related to fungal viruses (12), it is notable that Nackenback virus was found in both *Cx. pipiens* and *Cx. torrentium* libraries and was also more abundant than host non-RNA in the *Cx. torrentium* libraries (Table 2, Table 3). Hence, this virus may be truly mosquito-associated. We also detected two orthomyxoviruses, Wuhan Mosquito Virus 6 and Wuhan Mosquito Virus 4, both of which have previously been found in pools of *Culex* mosquitoes and are known to be mosquito associated (12).

### Double-stranded RNA viruses

We identified a total of eight double-stranded RNA viruses in our Swedish mosquitoes, all of which were novel. These belong to the *Partitiviridae* (N = 2), *Reoviridae* (N = 1) and *Toti/Chrysoviridae* (N = 5). For the family *Reoviridae*, Valmbacken virus clustered with *Aedes camptorhynchus* reo-like virus, previously discovered in mosquitoes (12). Valmbacken virus was also abundant and found in all libraries, and is therefore most likely a *Culex* associated reovirus (Table 2, Table 3).

In comparison, four of the five novel totiviruses clustered with other mosquito associated totiviruses (Figure 6), but only two (Lindangsbacken virus and Eskilstorp virus) were also found to be abundant. The fifth totivirus, Ahus virus, was highly divergent, had low abundance and clustered distantly with potentially protist originating viruses (Figure 6). Thus, the host association of Ahus virus remains uncertain.

The two novel partiti-like viruses, Vivastbo virus and Sonnbo virus, did not cluster with any viruses sequenced from mosquitoes, but rather grouped with viruses originating from various arthropod hosts (Figure 6). However, the relatively high abundance levels of Vivastbo virus suggest that it may be associated with mosquitoes (Table 2, Table 3).

Finally, all sequencing libraries generated here were negative for *Wolbachia* as assessed by mapping against the COX1 and WSP genes of *Wolbachia pipientis.* Although it is not possible to completely exclude the presence of other *Wolbachia* variants, our results suggest that differential presence/absence of *Wolbachia* has not affected the viral load or diversity observed.

## Discussion

Through total RNA-sequencing of 270 *Culex* mosquitoes collected in Sweden we identified 40 viruses, including 28 that are novel. A virome comparison between the two vector species *Cx. pipiens* and *Cx. torrentium* revealed that although these mosquitoes are from the same genus and have overlapping geographical distribution, the virome family and species composition and abundance differed to some extent between the two species, and also by the geographic location of sampling (Figure 1, Figure 2, Table 2).

Viewed at the family/order level, the relative virome abundance of *Cx. pipiens* was dominated by nido-, luteo-, and orthomyxo-viruses. In comparison, *Cx. torrentium* had a greater proportion of picorna-, nido-, mononega-, and reo-viruses. It should be noted, however, that family-wide comparisons could be significantly skewed in the presence of single highly abundant viruses, as was the case here (e.g. the Nam Dinh virus in library 2 that reached 30% of all reads in the library), such that analyses of relative abundance and diversity are better conducted at the species level. Viewed at the level of species per region, about one quarter of the viruses were unique to their respective sampling location (Figure 3). This suggests that local acquisition, as well as local ecosystem and habitat composition, may be important in shaping virome compositions.

Direct comparisons between virome studies are greatly complicated by such factors as differences in sequencing technologies, bioinformatic analyses, criteria for species demarcation, and study focus. Despite these important caveats, it is noteworthy that the number of viruses found in here is of a similar magnitude and diversity to those found at lower latitudes (12, 15). Hence, the virome composition appears not to follow the same trend as mosquito biodiversity, with fewer species in temperate regions (17, 18). Specifically, 24 different viruses were found in *Cx. torrentium*, of which six were unique to that species, and 34 viruses were found in *Cx. pipiens*, of which 16 were unique. Hence, 18 viruses were shared between both mosquito species, 16 of which we tentatively consider to be mosquito associated based on their abundance and phylogenetic position (Figure 3, Figure 4–6, Table 3).

Given their relatively close phylogenetic relationship (Figure 7) and the fact that both mosquito species inhabit the same region, share larval habitat (4), and blood-meal hosts (Hesson, J. unpublished), the difference in their viromes is striking. By considering virus abundance and phylogenetic position we suggest that 26 of the viruses discovered were likely mosquito-associated (Figure 4–6, Table 3), although we cannot exclude either false-negative or false-positive associations. For example, the divergent Ista virus (*Picornaviridae*) was found in high abundance and in multiple libraries, but did not cluster with any viruses that originated from mosquitoes, although it did group with other arthropods (Figure 4–6). The fact that it did not cluster with other mosquito viruses is, however, perhaps unsurprising as studies from temperate regions are few, and this is the first study investigating the virome of *Cx. torrentium*. It is clear that many viruses are seemingly ubiquitous in mosquitoes, covering a wide variety of climates and habitats (12, 15, 21), but whether Ista virus and many other viruses are truly mosquito-associated will need to be considered in additional studies. It was also noteworthy that no insect-specific flaviviruses were discovered in this study, even though these are relatively commonplace (22) and have previously been found in mosquitoes in northern Europe (23, 24).

**Figure 7.**
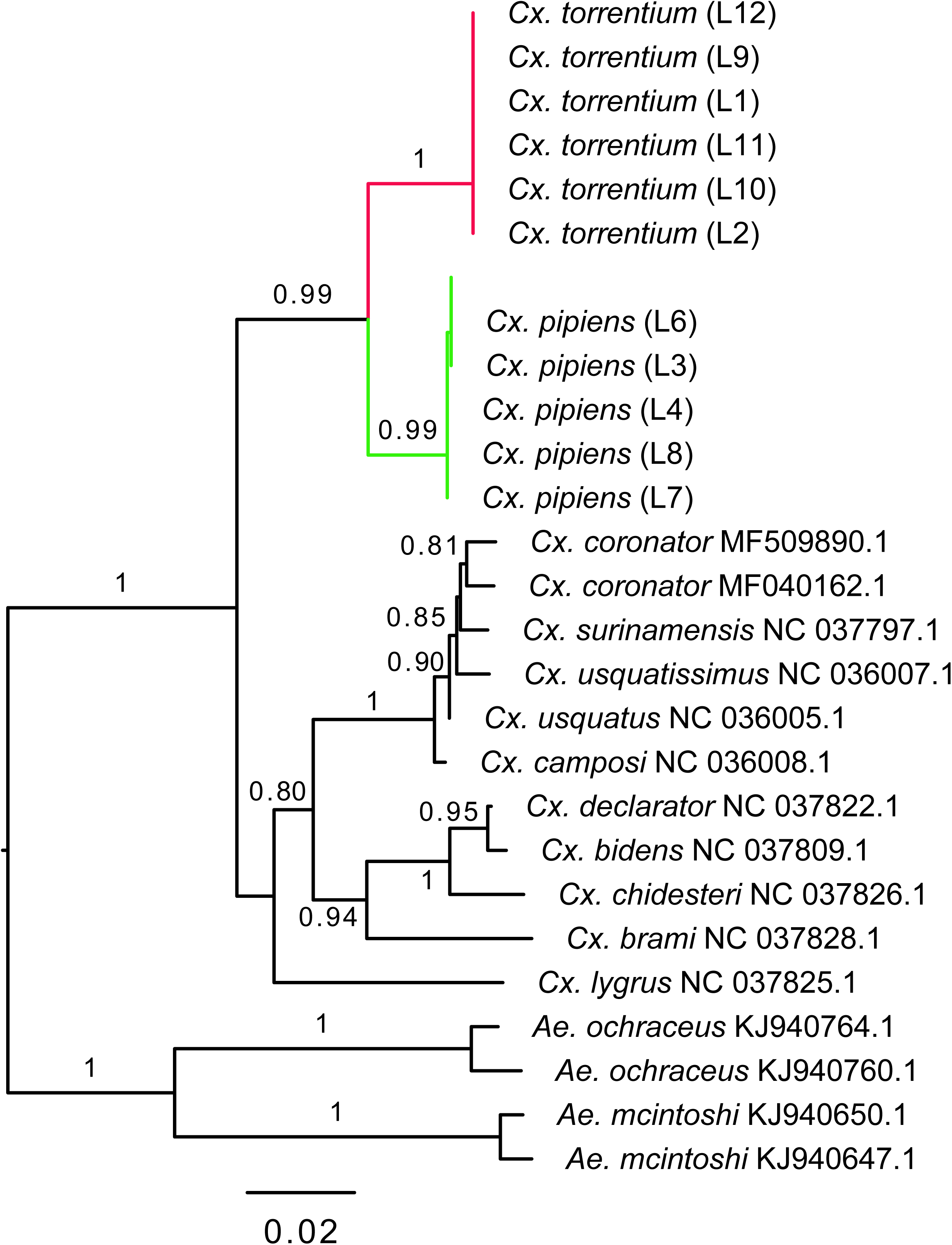
Phylogenetic relationships, based on partial COX1-gene, of *Cx. pipiens* and *Cx torrentium* for all libraries (L1–L12) together with representative publicly available reference sequences (with their associated GenBank accession numbers). Numbers on branches indicate SH support, and only branches with SH support ≥80% are indicated. Branch lengths are scaled according to the number of nucleotide substitutions per site. The tree is midpoint rooted for clarity only.

Notably, our study suggests that pathogenic viruses, such as SINV, can sometimes have similar abundance to viruses not associated with human disease (Table 2), suggesting that presence of pathogenic viruses does not necessarily impact overall virome composition. The difference in viral load of certain viruses between *Cx. torrentium* and *Cx. pipiens* is interesting. One potential explanation is that the virome composition has been impacted through differential associations with the intracellular bacteria *Wolbachia pipientis*. Indeed, it has previously been shown that *Cx. pipiens* is commonly infected with *Wolbachia*, while this bacterium is absent from *Cx. torrentium* (9). Wolbachia is well-known for its ability to influence virus infection in mosquitoes, but has mostly been studied in systems with pathogenic viruses such as dengue (25). However, we found no compelling evidence for *Wolbachia* in any of the *Culex* samples studied here.

The species separation between *Cx. pipiens* and *Cx. torrentium* has long been ignored, largely because of the need for molecular demarcation, so that it has been assumed that most of the biology of the two species is comparable. Our study reveals that *Culex* mosquitoes in northern temperate regions harbour as much, and sometimes more, viral diversity as mosquitoes in tropical and sub-tropical regions. We also show that the two vector species share many viruses, but also differ noticeably in which RNA viruses they harbour, and at significantly different abundance levels. Further studies should aim to study how mosquito host structure and feeding preference, evolutionary history, ecological relationships, geography and environment determines virome composition and the potential interaction with pathogenic and non-pathogenic viruses, as well as the impact of virome composition on mosquito health.

## Materials and Methods

### Mosquito collection

Mosquitoes were collected from two regions in Sweden: (i) from floodplains of the Dalälven river in central Sweden (60.2888; 16.8938) in 2006, 2009, and 2011, and (ii) around the city of Kristianstad, in southern Sweden (56.0387; 14.1438) in 2006 and 2007. Collections were performed using CDC-light traps baited with carbon dioxide, and catches were sorted and identified to species on a chilled table, using keys by Becker et al. (26). In total, legs from 270 *Cx. pipiens/torrentium* mosquitoes were removed to enable molecular identification to species (27). Bodies were homogenized in PBS buffer supplemented with 20% FCS and antibiotics and stored at −80C degrees until further processing.

### Sample processing and sequencing

Total RNA was extracted from 12 pools from the homogenate of individual *Cx. torrentium* (n = 150) and *Cx. pipiens* mosquitoes (n = 120) (Supplementary table 1), using the RNeasy^®^ Plus Universal kit (Qiagen) following the manufacturer’s instructions. The extracted RNA was subsequently DNased and purified using the NucleoSpin RNA Clean-up XS kit (Macherey-Nagel). Prior to library construction, ribosomal RNA (rRNA) was depleted from the purified total RNA using the Ribo-Zero Gold (human-mouse-rat) kit (Illumina) following the manufacturer’s instructions. Sequencing libraries were then constructed for all rRNA-depleted RNA-samples using the TruSeq total RNA library reparation protocol (Illumina). All libraries were sequenced on a single lane (paired-end, 150 bp read-length) on an Illumina HiSeq X10 platform. Library preparation and sequencing was carried out by the Beijing Genomics Institute (www.bgi.com/global/). All 12 libraries were quality trimmed with trimmomatic v.0.36 (28) and then assembled *de novo* using Trinity v.2.5.4 (29).

### Discovery of viruses and Wolbachia bacteria

Trinity assemblies were screened against the complete non-redundant NCBI GenBank nucleotide (nt) and protein (nr) databases using blastn and diamond (30) blastx with a cut-off e-value of 1×10^−5^. Assemblies identified as RNA viruses were screened against the Conserved Doman Database (www.ncbi.nlm.nih.gov/Structure/cdd/wrpsb.cgi) with an expected value threshold of 1×10^−3^ to identify viral sequence motifs. The mitochondrial COX1 gene, mined from the sequence data, and all contigs with RdRp-motifs was mapped back, using Bowtie2 (31), against all quality trimmed libraries to estimate abundance. A virus was considered to be in high abundance if: (i) it represented >0.1% of total non-ribosomal RNA in the library, and (ii) if the abundance was higher to that of abundant host COX1 gene (12, 32), and hence likely to be mosquito associated. Hits that were below the level of cross-library contamination due to index-hopping, measured as 0.1% of the most abundant library for the respective virus species or less than 1 read per million mapped to a specific virus contig, was considered negative (coloured grey in Table 1 and Table 2, respectively). To investigate the presence of *Wolbachia* bacteria in the libraries, published sequences of the *Wolbachia Cx. pipiens* wsp surface protein gene (DQ900650.1) and the mitochondrial COX1 gene (AM999887.1) were mapped backed against all libraries using the above criteria for abundance and presence/absence.

### Inference of virus evolutionary history and host associations

The evolutionary (i.e. phylogenetic) history of the viruses discovered was inferred by aligning protein translated open reading frames with representative sequences from the *Alphaviridae*, (Order) *Bunyavirales, Endornaviridae, Luteoviridae*, (Order) *Mononegavirales*, Nido-like viruses, (Order) *Orthomyxovirales, Partitiviridae, Picornaviridae*, Qin-like viruses, *Reoviridae, Totiviridae, Tymoviridae* and *Virgaviridae* and Negev-like viruses. All RdRp amino acid sequence alignments were performed using the E-INS-i algorithm in Mafft (33). Poorly aligned regions, in which amino acid positional homology could not be confirmed, were then removed from the alignments using TrimAl utilizing the ‘strict’ settings. Finally, phylogenetic trees were computed with a maximum likelihood approach as implemented in PhyML (34) employing the LG +Γ amino acid model, SPR branch-swapping and the approximate likelihood ratio test (aLRT) with the Shimodaira-Hasegawa-like procedure used to assess branch support. The resultant phylogenetic trees were edited and visualised with FigTree v.1.4.2 (http://tree.bio.ed.ac.uk/so ware/figtree).

To help assess whether the novel viruses discovered are mosquito-associated – that is, to distinguish those that actively replicate in the host from those present in diet mosquito or a co-infecting micro-organism – we considered four factors: (i) the abundance of viral contigs per total number of reads in a library, (ii) the abundance in relation to the COX1 host gene, (iii) the presence in the libraries, and (iv) phylogenetic clustering with other mosquito derived viruses.

The raw sequence data generated here has been deposited in the NCBI short read archive (BioProject: PRJNA516782) and all viral contigs has been deposited in NCBI GenBank (accession numbers: MK440619–MK440659).

## Acknowledgments

The authors are grateful to Dr. JO Lundström for access to mosquito material. JHOP is supported by the Swedish research council FORMAS (grant nr: 2015-710). ECH is supported by an ARC Australian Laureate Fellowship (FL170100022). JCH is supported by E and R Börjeson’s Foundation and the Swedish Society for Medical Research.

## Supplementary Material

**Supplementary Table 1.** Information on collection site, year of collection and pool size for all *Cx. pipiens* and *Cx. torrentium* libraries.

**Supplementary Table 2.** Number of reads mapped to each virus, *Wolbachia* and host genes per library per mosquito species.

